## Supplementary Figures for "Imbalanced segregation of recombinant haplotypes in hybrid populations reveals inter- and intrachromosomal Dobzhansky-Muller incompatibilities"

A

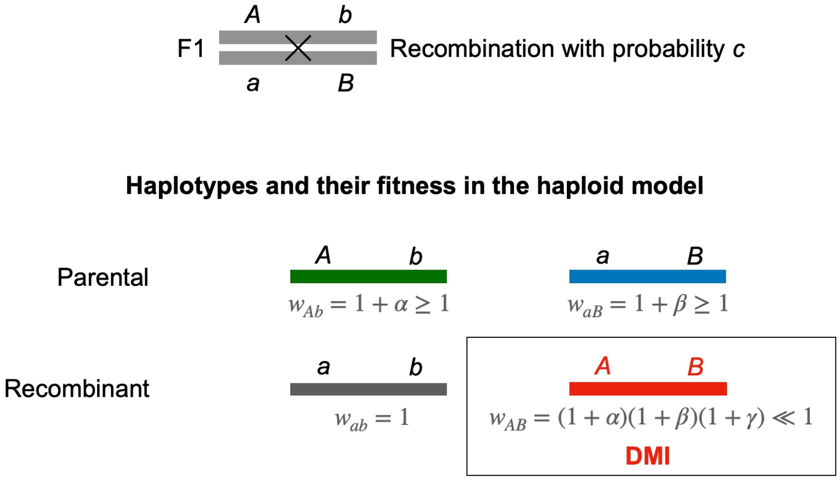


B

**Selection coefficients with a DMI vs. HRI**

| | *ab* | *Ab* | *aB* | *AB* |
|---|---|---|---|---|
| DMI | – | ↑ | ↑ | ↓↓ |
| | 1 | 1.001 | 1.002 | ≈0.5 |
| HRI | – | ↓ | ↓ | ↓↓ |
| | 1 | 0.9 | 0.89 | ≈0.8 |

Figure S1. Illustration of the genomic composition and the fitnesses in the haploid model. A. In our toy model of hybridization with a single DMI pair, two recombinant haplotypes (*ab* and *AB*) are generated from the parental haplotypes (*Ab* and *aB*) from the F2 onwards. In the following generations, all four haplotypes are segregating. In the Dobzhansky-Muller incompatibility (DMI) model, the recombinant haplotype *AB* is strongly selected against. Throughout the paper we allow for weak direct selection for the derived alleles (here, $\alpha$ = 0.001 for *A* and $\beta$ = 0.002 for *B*) and an intermediate to strong fitness interaction between *A* and *B* (here, $\gamma$ = -0.5). B. Recombinant imbalance caused by a DMI as compared with Hill-Robertson interference (HRI). Alleles *A* and *B* under strong selection in the same direction can lead to recombinant imbalance between *ab* and *AB*. The arrows show the intensity of the absolute reduction of each haplotype frequency in one generation.

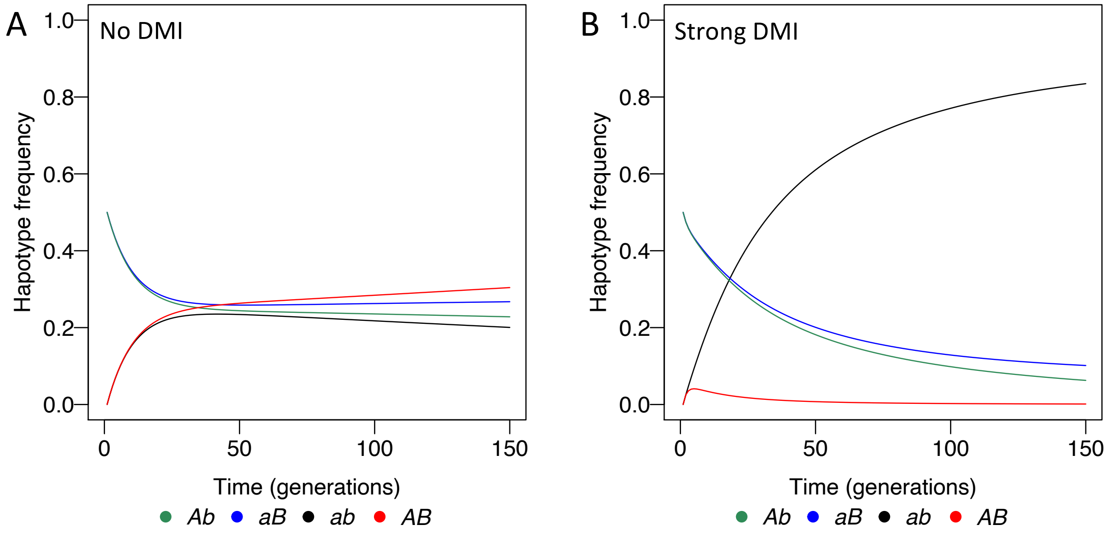


Figure S2. Deterministic frequency dynamics of the four haplotypes in a new hybrid population in the deterministic haploid model. Alleles *A* and *B* are under weak direct selection ($\alpha = 0.001$ for *A* and $\beta = 0.002$ for *B*). The epistatic interaction between alleles *A* and *B* is $\gamma$. The recombination probability $c$ is 0.1. The admixture proportion is 0.5. A. The haplotype frequencies approach linkage equilibrium soon after hybridization without a DMI ($\gamma = 0$). B. With a strong DMI ($\gamma = -0.5$) the emerging recombinant haplotype *AB* is eliminated quickly, whereas the *ab* haplotype has a marginal advantage that drives it to high frequencies.

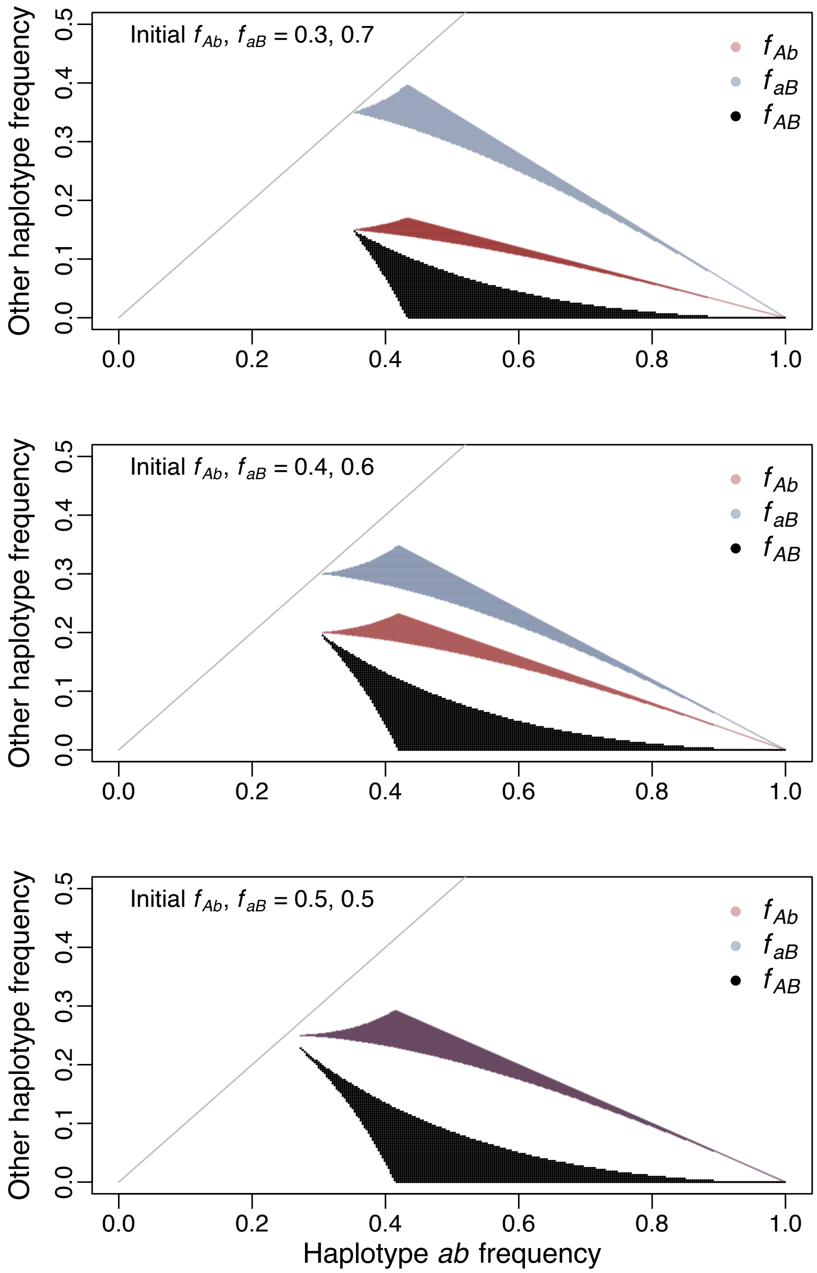


Figure S3. Haplotype frequencies that create negative $X(2)$. The three panels represent three different admixture proportions for *Ab* and *aB*. The frequencies of four haplotypes $(f_{aB}, f_{Ab}, f_{ab}, f_{AB})$ were plotted. Haplotype frequency of *ab* $(f_{ab})$ is shown on the x axis and the frequencies of the other three haplotypes are shown on the y axis. The ratio of $f_{aB}$ and $f_{Ab}$ stays the same as their initial ratio for every combination of haplotype frequencies $(f_{aB}, f_{Ab}, f_{ab}, f_{AB})$. The slope and intercept of the grey diagonal line are 1 and (0,0). As the combination of four haplotype frequencies always lie under the grey line, the haplotype *ab* has always the largest frequency. The haplotype *AB* has always the smallest frequency. In this figure, $f_{ab}$ could be close to but smaller than 1.

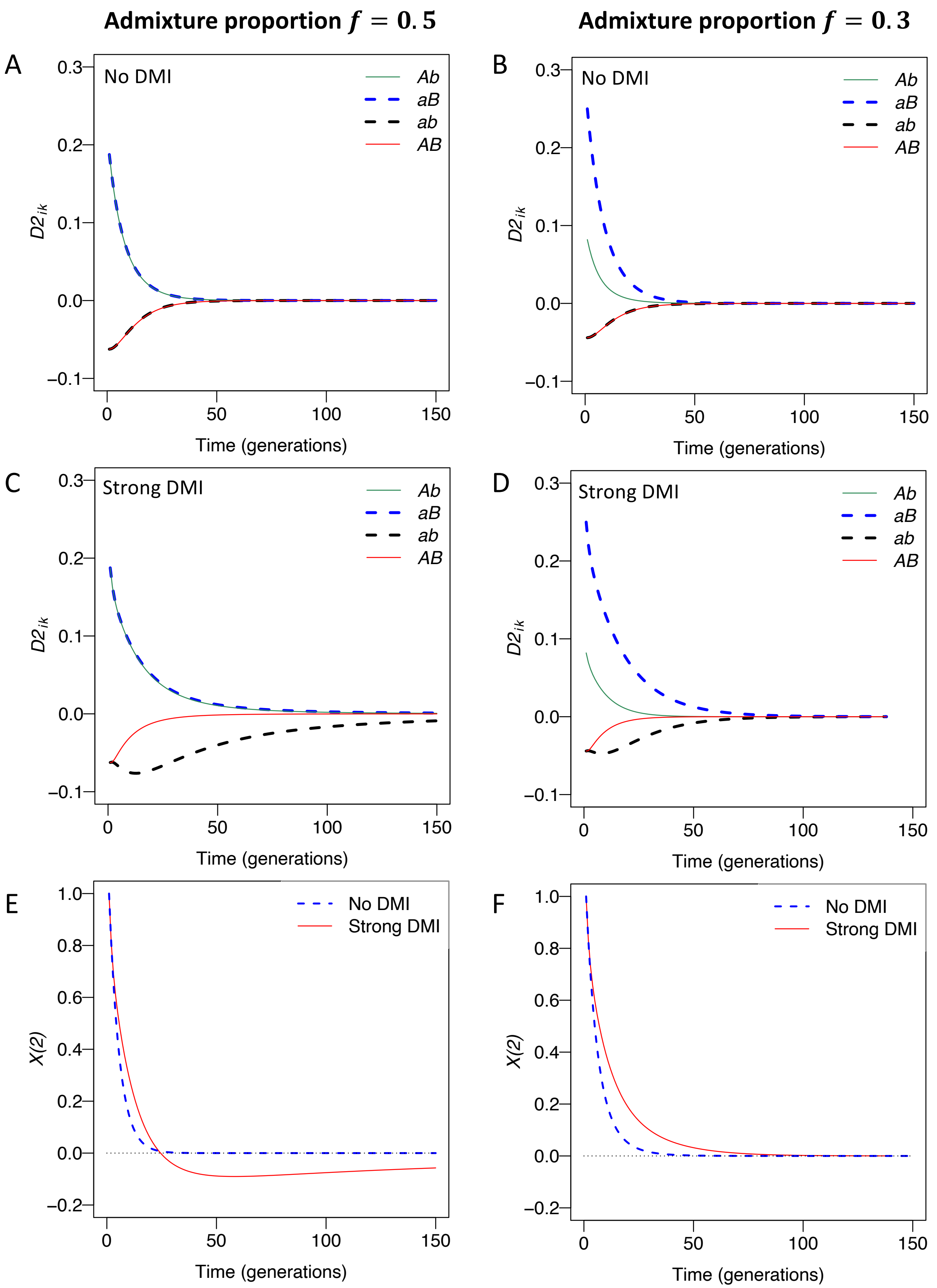


Figure S4. Dynamics of $D2_{ik}$ and $X(2)$. A-D. $D2_{ik}$ in four scenarios combining equal/unequal admixture proportions and with/without DMIs, obtained from the deterministic haploid model. E-F. $X(2)$ trajectory with (blue dashed line) and without (red line) DMIs. The direct selection coefficients are $\alpha$= 0.001, $\beta$ = 0.002. The strength of epistasis is $\gamma$ = -0.5 (strong DMI) and $\gamma$ = 0 (no DMI). The recombination probability is 0.1.

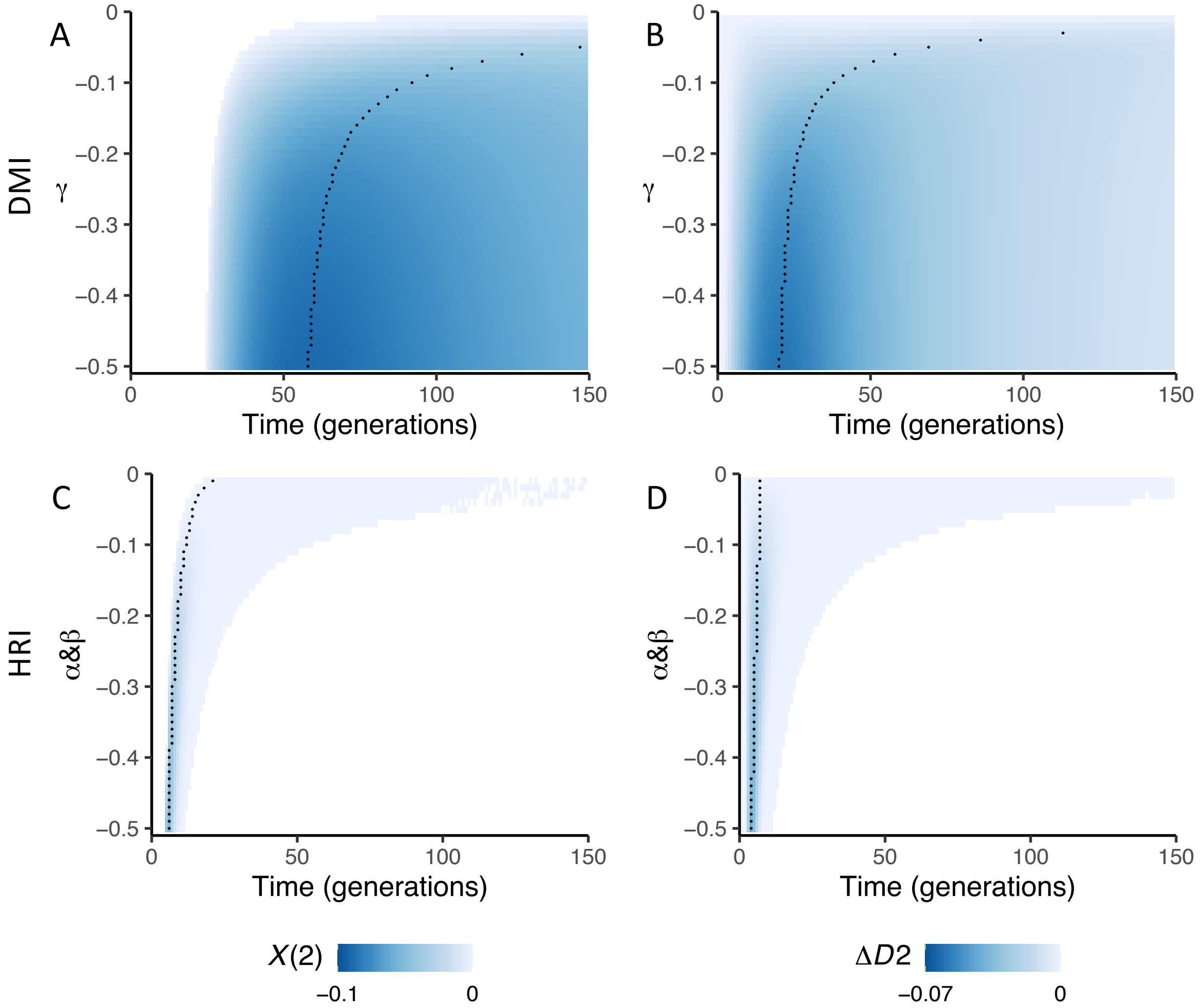


Figure S5. Comparison of the dynamics of $X(2)$ and $\Delta D2 = D2_{ab} - D2_{AB}$ for a DMI (A&B) compared with strong direct selection (Hill-Roberson interference, HRI, C&D) in the deterministic haploid model. The direct selection coefficients acting on $A$ and $B$ are $\alpha, \beta$. The admixture proportion is $f = 0.5$. Black dots highlight the generation with the largest $-X(2)$ or $-\Delta D2$ for each trajectory. The values of $X(2)$ and $\Delta D2$ are represented by blue color intensity. Blank areas represent either $X(2)$>0 (to the left of the blue area) or at least one locus close to fixation (minor allele frequency <0.005, on the right of the blue area). The recombination probability between the two DMI loci is 0.1. A&B. DMIs ($\alpha = 0.001, \beta = 0.002,\ \gamma < 0$ varies on the y axis) exist for a longer longer time in the population than HRI (C&D). C&D. HRI ($\gamma = 0, \alpha\&\beta < 0$, y axis) can briefly induce weak negative $X(2)$ and $\Delta D2_{ik}$ earlier than a DMI, but the two-locus polymorphism is quickly purged from the population.

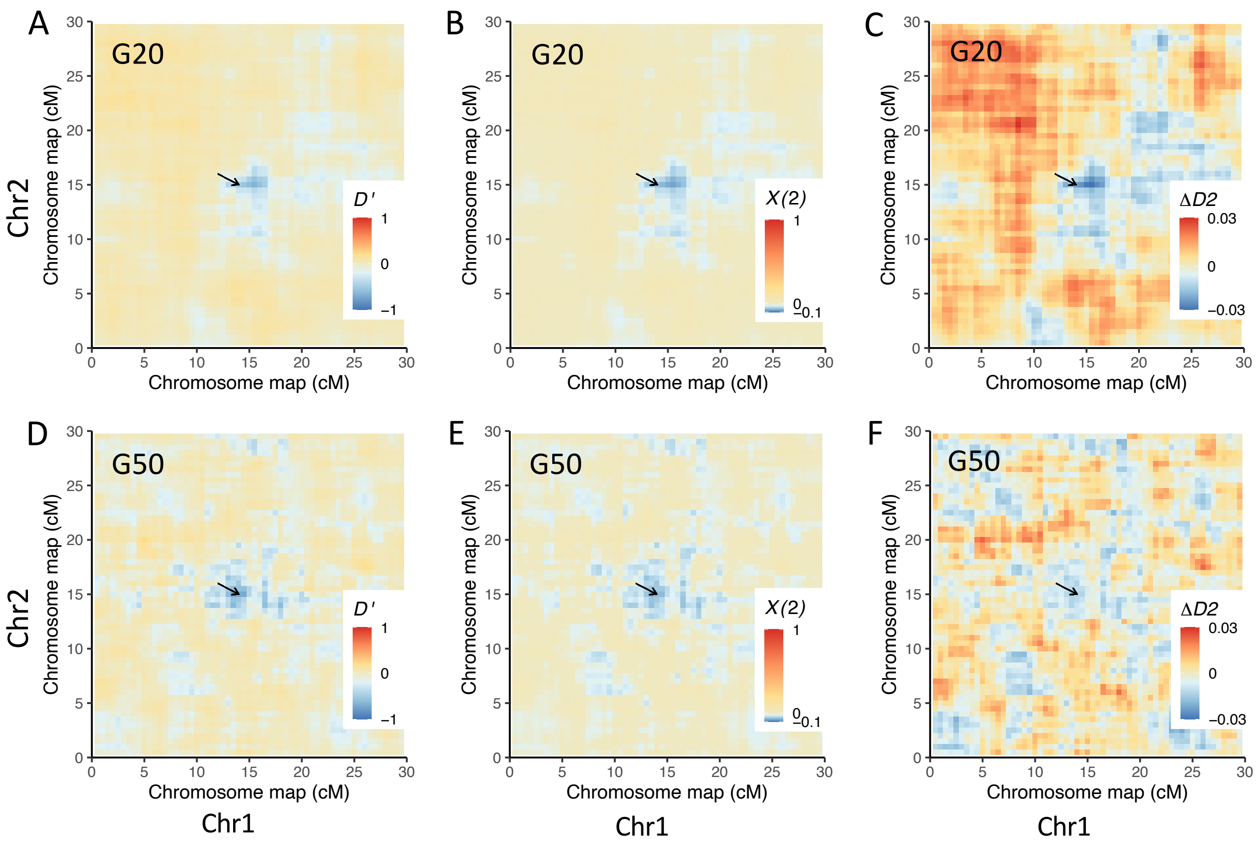


Figure S6. Negative $X(2)$ pinpoints an intrachromosomal DMI in simulated data. The distribution of $X(2)$, $D'$, and $\Delta D2$ of a chromosome with a DMI pair at generation 50 under a Wright-Fisher model for one simulation run using SLiM. The direct selection coefficients are $\alpha = 0.001, \beta = 0.002$. The strength of epistasis is $\gamma = -0.5$. The interacting loci generating the DMI were located at chromosome 1 (chr1):15 cM and chr2: 15 cM, such that the recombination probability was $c = 0.5$ between the two loci. A-C. Heatmaps of $D'$, $X(2)$ and the $\Delta D2$ within the simulated chromosome at generation 20 (G20). D-F. Heatmaps of $D'$, $X(2)$ and the $\Delta D2$ within the simulated chromosome at generation 50 (G50). Only the DMI loci and their 10 cM flanking regions were shown in D-F. Black arrows highlight the location of the DMI.

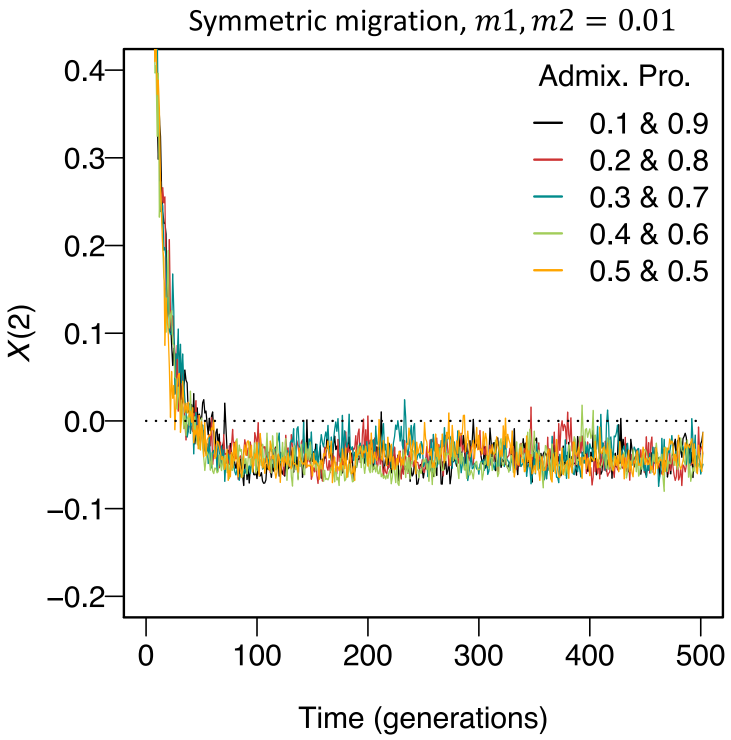


Figure S7. Negative X(2) dynamics a DMI persist for long times if there is ongoing immigration from both parental populations. Each line is one simulation run using SLiM, assuming a strong DMI ($\alpha = 0.001, \beta = 0.002, \gamma = -0.5$) under a Wright-Fisher model. The recombination probability between the interacting loci was 0.1. 300 individuals were sampled in each simulation. Trajectories were plotted from generation 2 onwards. DMIs can be maintained by symmetric migration from two parental populations ($m1, m2 = 0.01$) for a long time, and not affected by the initial admixture proportions (Admix. Pro.).

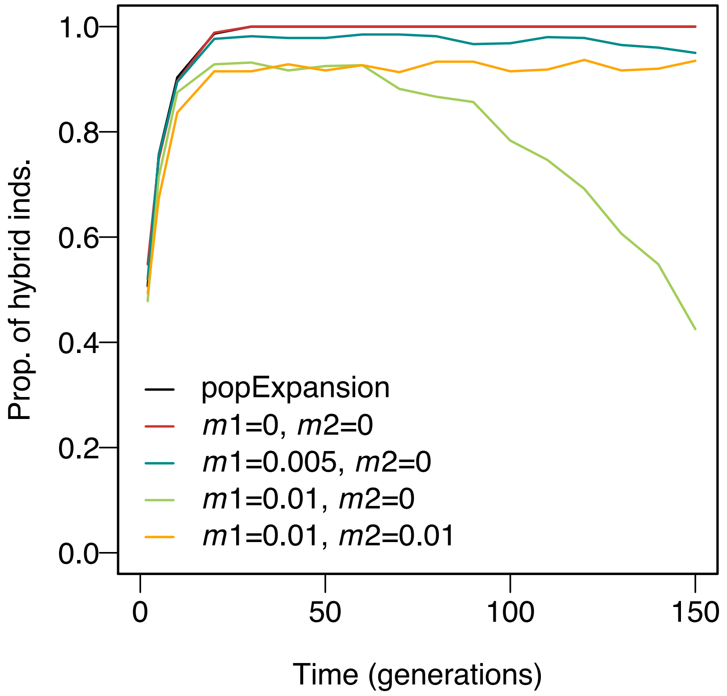


Figure S8. Proportion of hybrid individuals under five demographic scenarios. Here, hybrids are defined as individuals that derived at least 10% of their genome from each parental population. Each line is one simulation run using SLiM, assuming equal admixture proportions of the parental populations and a strong DMI ($\alpha = 0.001, \beta = 0.002, \gamma = -0.5$) under a Wright-Fisher model. The recombination probability between the interacting loci was 0.1. Three hundred individuals were sampled in each simulation.

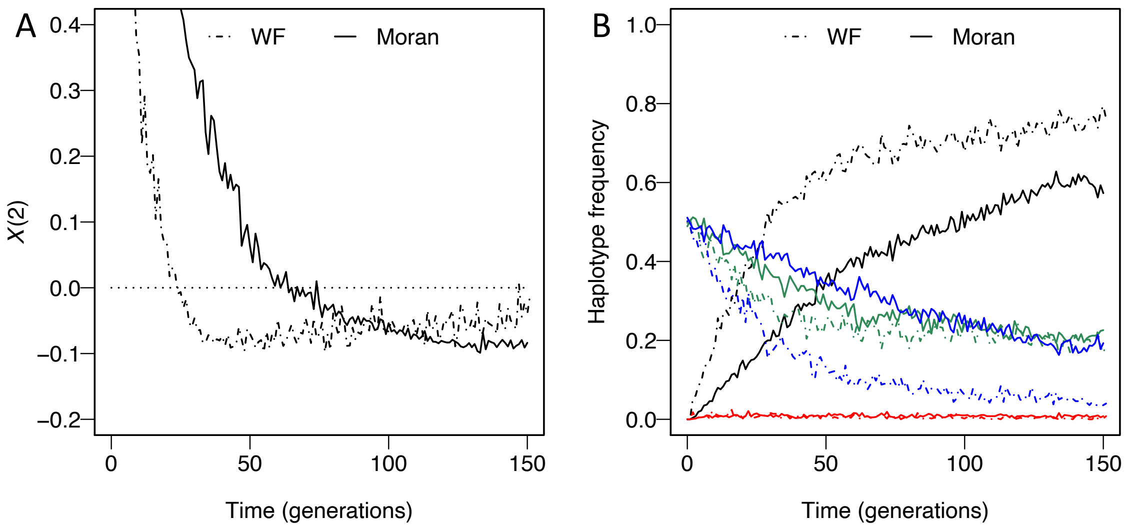


Figure S9. Dynamics of haplotype frequencies and $X(2)$ for a DMI under diploid Wright-Fisher (WF; dash-dotted line) and Moran (solid line) models for one simulation run using SLiM, assuming a population size of 5000. The model parameters were $\alpha = 0.001, \beta = 0.002, \gamma = -0.5$, with recombination probability $c = 0.1$ between the DMI loci. Three hundred individuals were sampled in each simulation. A. Negative $X(2)$ appears ~50 generations earlier under a Wright-Fisher model than under a Moran model. B. Haplotype frequencies change more quickly in the Wright-Fisher model than that in the Moran model. Lines indicate the haplotype frequencies of *AB* (red), *ab* (black), *Ab* (blue) and *aB* (green).

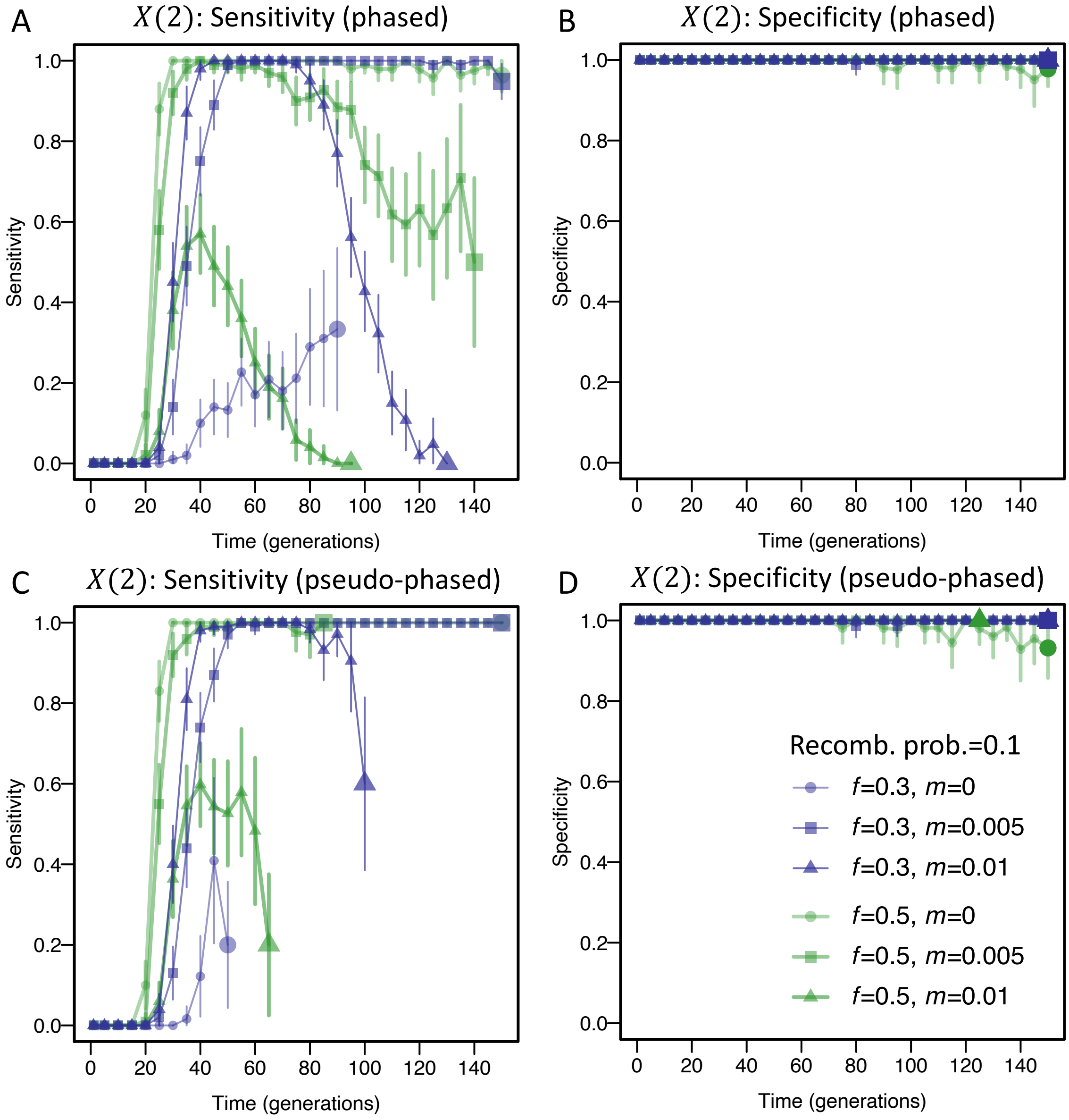


Figure S10. Sensitivity and specificity of negative $X(2)$ for detecting an intragenic DMI with c=0.1, computed from 100 simulations of each parameter combination using SLiM, assuming a Wright-Fisher population of 5000 individuals. We considered two admixture proportions, $f = 0.5$ and $f = 0.3$, and three migration rates: 0, 0.01, and 0.05. A strong DMI ($\alpha = 0.001, \beta = 0.002, \gamma = -0.5$) and no DMI ($\alpha = 0.001, \beta = 0.002,\ \gamma = 0$) were simulated under a Wright-Fisher model. The population size was 5000. Here, any interaction with $X(2) < -0.005$ & $D' < 0$ was taken as a putative DMI. Trajectories ended (indicated by large plot markers) when one of the interacting loci became monomorphic (A&B) or there is <50 homozygous genotypes for statistics (C&D). 300 individuals were sampled in each simulation. Vertical lines indicate the 95% confidence interval (Wald method). A&B Sensitivity and specificity using phased data. C & D. Sensitivity and specificity when only homozygous genotypes were used at each locus. A&C. Sensitivity is high unless strong migration leads to swamping of the DMI alleles. B&D. Specificity is high under all tested scenarios. Vertical lines are 95% confidence interval (Wald test).

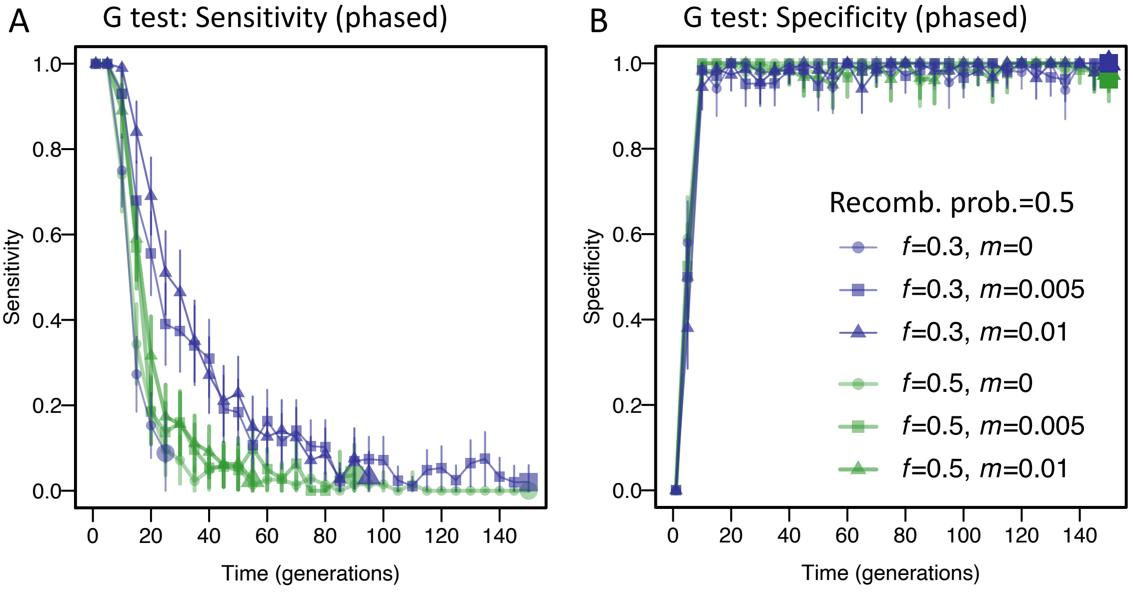


Figure S11. Sensitivity and specificity of the G test that indicates DMIs ($p < 0.05$). We considered two admixture proportions, $f = 0.5$ and $f = 0.3$, and three migration rates: 0, 0.005, and 0.01. A strong DMI ($\alpha = 0.001, \beta = 0.002, \gamma = -0.5$) and no DMI ($\alpha = 0.001, \beta = 0.002, \gamma = 0$) were simulated under a diploid Wright-Fisher model. The recombination probability was 0.5. For each parameter combination, we ran 100 simulations with SLiM. Sensitivity (A) was the proportion of successful DMI detections with the strong DMI. Specificity (B) was the proportion of true negative detections without DMI. Trajectories ended when one of the interacting loci became monomorphic or at generation 150 (indicated by large markers). Vertical lines are 95% confidence interval (Wald test).

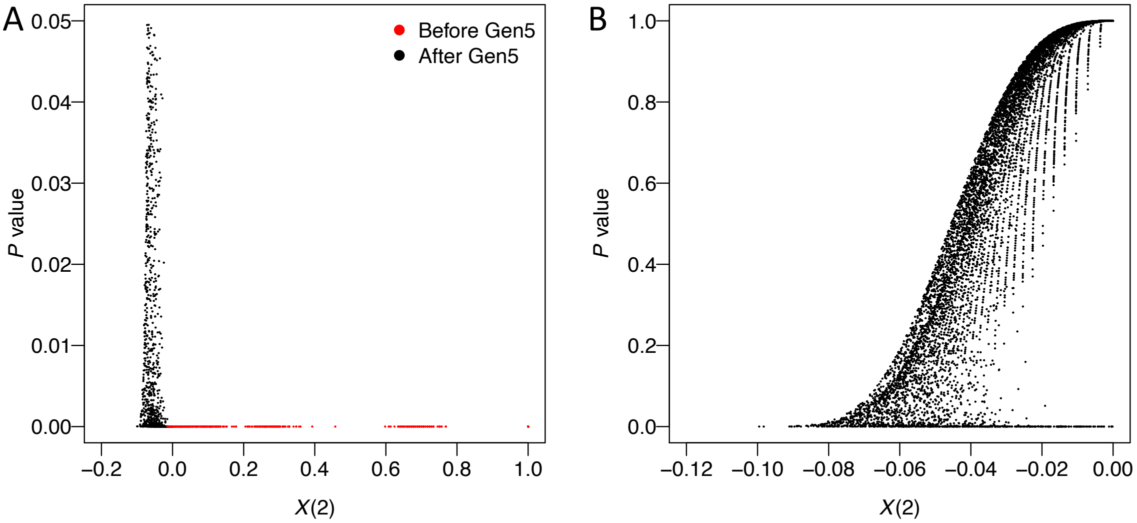


Figure S12. The distribution of $X(2)$ plotted against p values from the G test. Data were obtained from 100 simulations of a single DMI under a diploid Wright-Fisher model. Here, we used the same data as in Figure 5 and S11 with equal admixture proportions and no migration within 150 generations, i.e., the data points represent the $X(2)$ and G test statistics at the DMI loci, pooled across all sampling points. A strong DMI ($\alpha = 0.001, \beta = 0.002, \gamma = -0.5$) was simulated under a diploid Wright-Fisher model. The recombination probability was 0.5. The distribution was zoomed into two areas by two independent criteria for detecting DMI: (A) small p values by G test ($p < 0.05$); (B) negative $X(2)$ values. Red dots indicate DMIs that are observed before Generation 5 with a significant p value by the G test in panel A. Black dots indicate DMIs after Generation 5 in panel A and B. Only 9.9% of the simulations showed a significant p value by the G test ($p < 0.05$) in panel B.

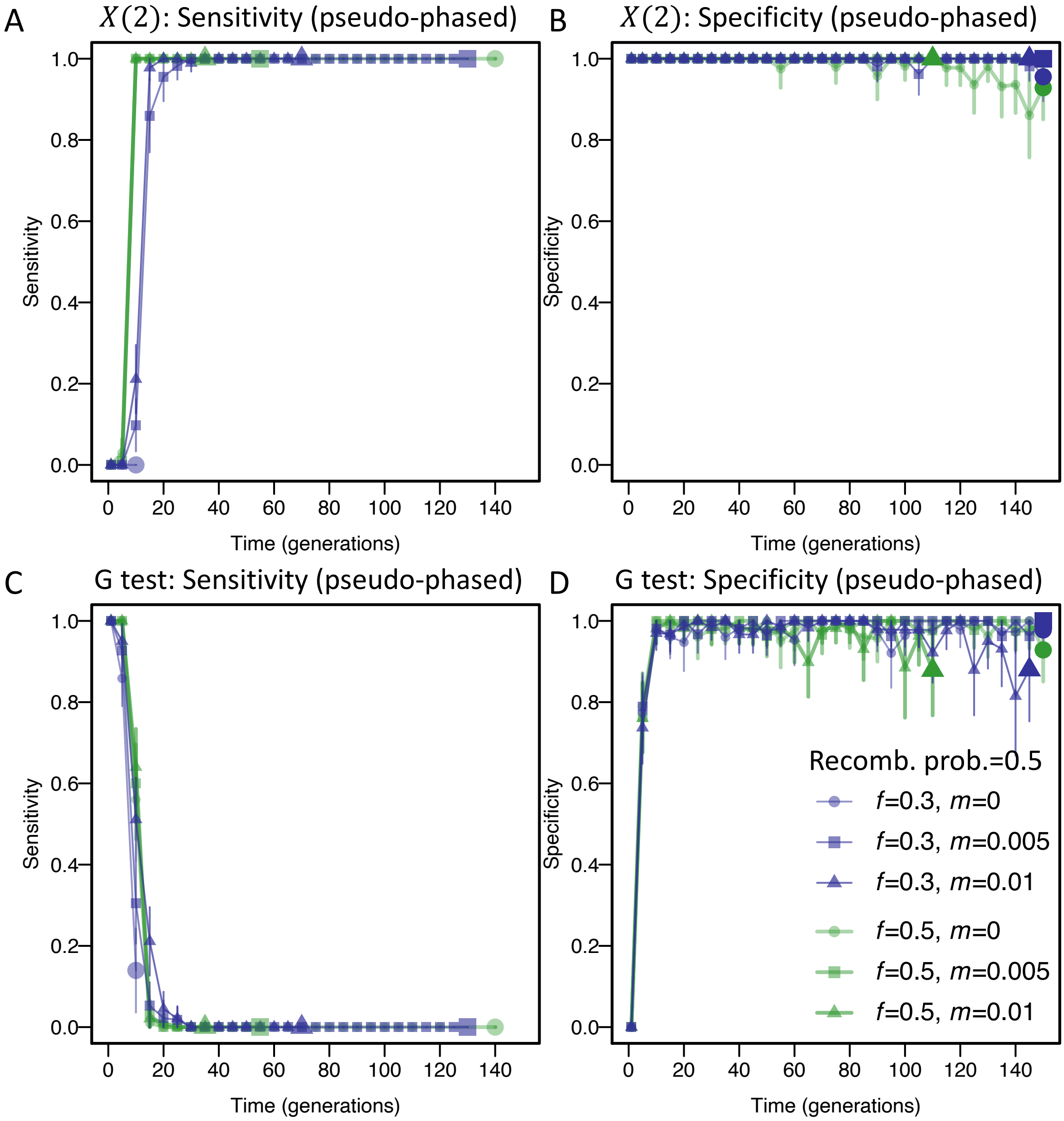


Figure S13. Sensitivity and specificity of negative $X(2)$ (A&B) and G test (C&D) indicating DMIs when only homozygous genotypes were used at each locus. We considered two admixture proportions, $f = 0.5$ and $f = 0.3$ and three migration rates ($m$): 0, 0.005 and 0.01. A strong DMI ($\alpha = 0.001, \beta = 0.002, \gamma = -0.5$) for sensitivity and no DMI ($\alpha = 0.001, \beta = 0.002, \gamma = 0$) for specificity were simulated under a diploid Wright-Fisher model. The recombination probability was 0.5. For each parameter combination, we ran 100 simulations using SLiM. In panel A&B, any interaction with $X(2) < -0.005$ & $D' < 0$ was taken as a putative DMI. In panel C&D, any interaction with $p<0.05$ was taken as a putative DMI. Sensitivity (A&C) was the proportion of successful DMI detections with the strong DMI. Specificity (B&D) was the proportion of true negative detections without DMI. Trajectories end when there is <50 homozygous genotypes for statistics or at generation 150 (indicated by large markers). Vertical lines are 95% confidence interval (Wald test).

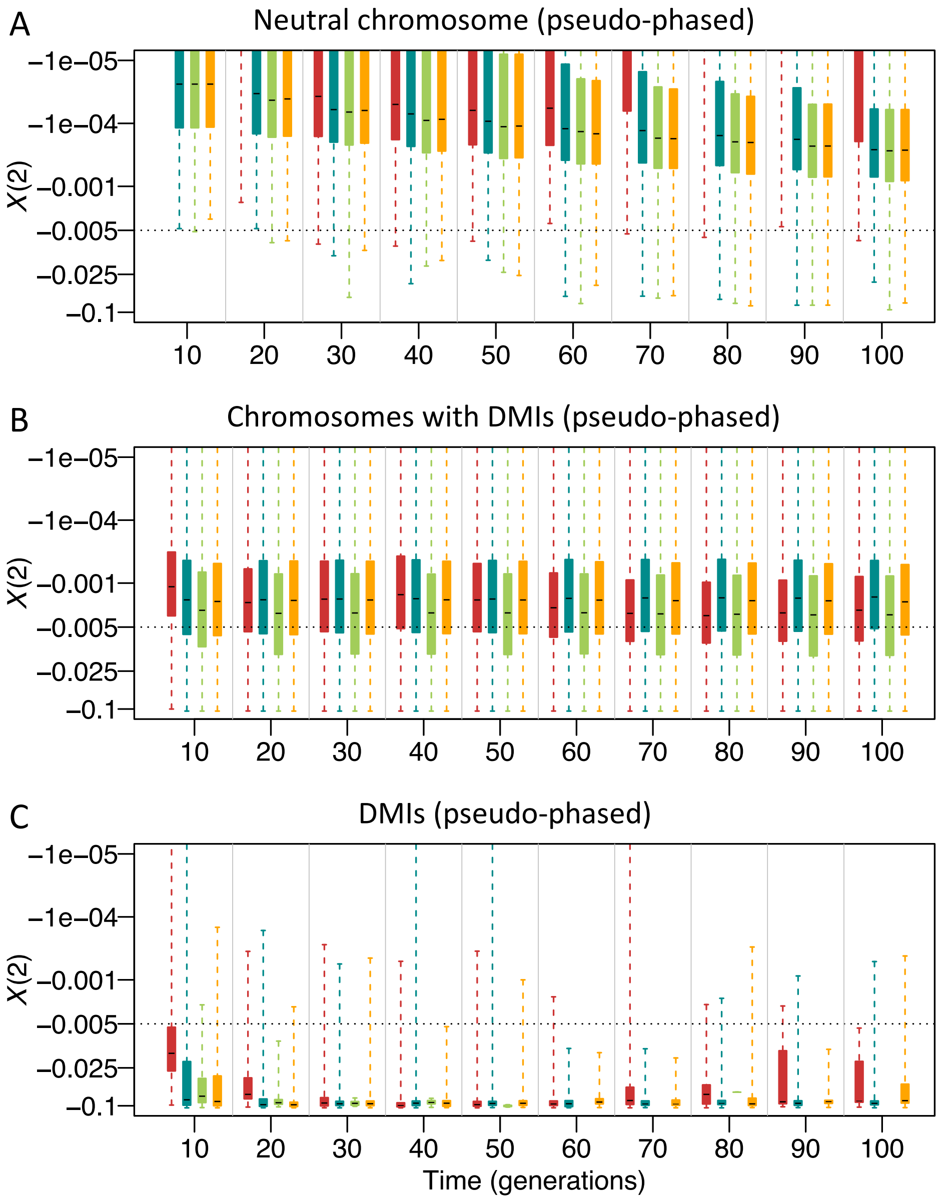


Figure S14. Distributions of negative $X(2)$ when $D' < 0$ for DMI detection under four demographic scenarios. This figure represents two starting frequencies of the minor parental population, $f = 0.5$ and $f = 0.3,$ and different immigration rates from the two parental populations ($m1$ from minor parental population, $m2$ from major parental population). Light green boxes show the $X(2)$ distribution for the three-locus two-DMI model (3-locus) and the other boxes for the four-locus two-DMI model. DMIs are randomly distributed on four chromosomes. Direct selection coefficients on two incompatible alleles were drawn from an exponential distribution with mean = 0.001. Epistatic coefficients were drawn from a uniform distribution [-1, -0.001]. We calculated $X(2)$ statistics using homozygous genotypes from a sample of two hundred individuals for each simulation. We simulated each scenario for 100 times with SLiM. $X(2)$ values were summarized for three types of genomic regions, neutral chromosomes (A), chromosomes with DMIs (B), and individual DMI loci (C). Boxplots represent the interquartile range; whiskers extend to minimal and maximal values. The dash line is $X(2) = -0.005$.

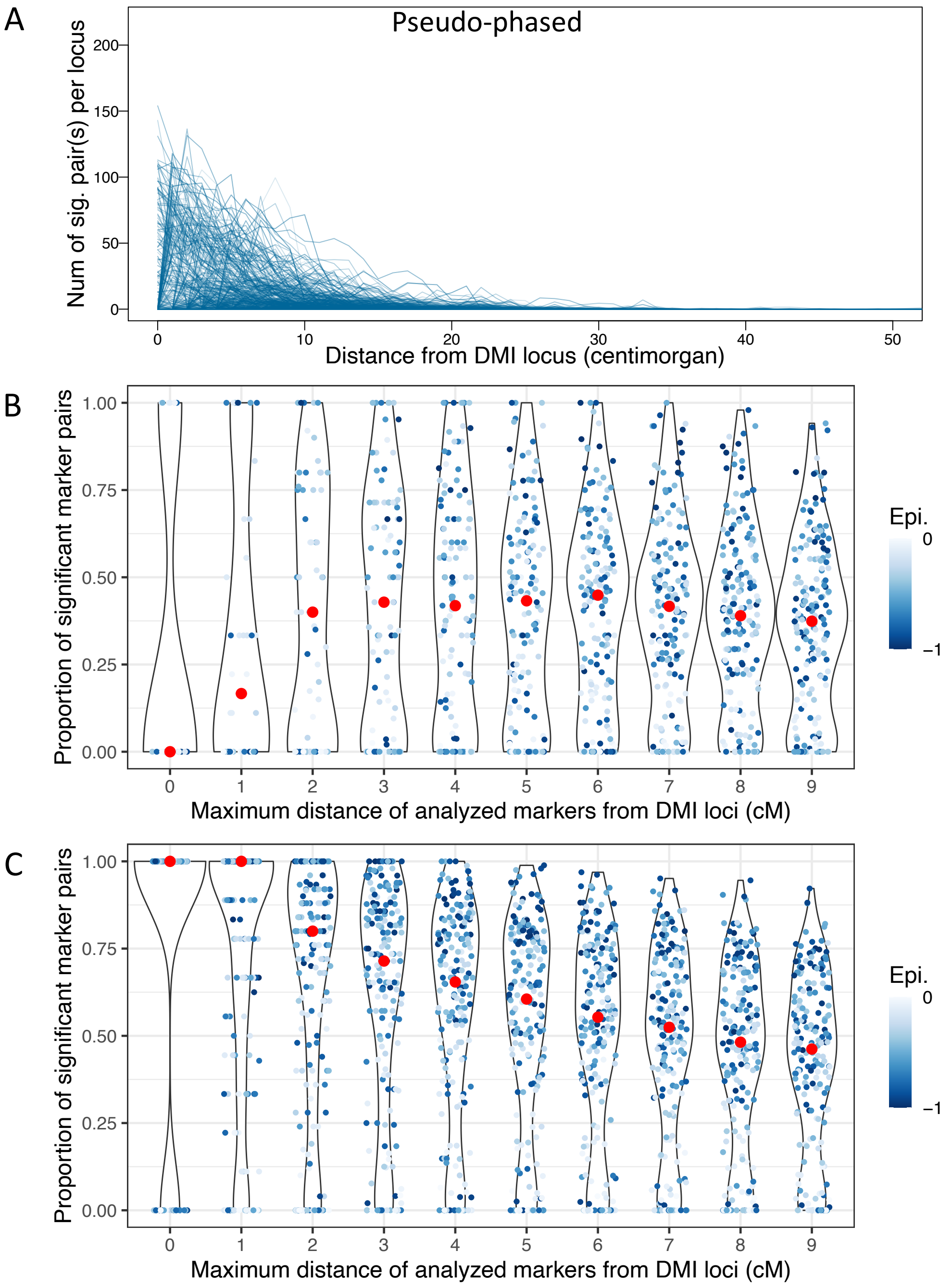


Figure S15. Propagation of the DMI signal into flanking regions in simulated data (sampled at generation 30, significance threshold: X(2)<-0.005 & D'<0) when only homozygous genotypes were sampled (sampling criteria for panels A&B: homozygous genotype frequency >0.05, heterozygosity<0.6; panel C shows homozygous data without further filtering). Data resulted from 100 simulations using SLiM, in which two DMIs (i.e., four DMI loci) were randomly distributed on four chromosomes, thus resulting in inter- and intrachromosomal DMIs. The direct selection coefficients α and β were drawn from exponential distribution with mean value 0.001. The strength of epistasis was drawn from a uniform distribution, γ∈[-1, -0.001], which is represented by the blue colour intensity in panels B&C. To resemble the demography of the studied hybrid swordtail fish populations, the admixture proportion was set to f=0.3 and the migration rate was m=0.01. 200 individuals were sampled from each simulation to obtain the statistics. A. The number of significant marker pairs decreases rapidly with distance to the nearest true DMI locus. B&C. Proportion of significant marker pairs when statistics were computed for all pairs between two increasingly large regions around the true interacting DMI loci, with (B) and without (C) strict sampling criteria. Red dots indicate the median obtained from 100 simulations.

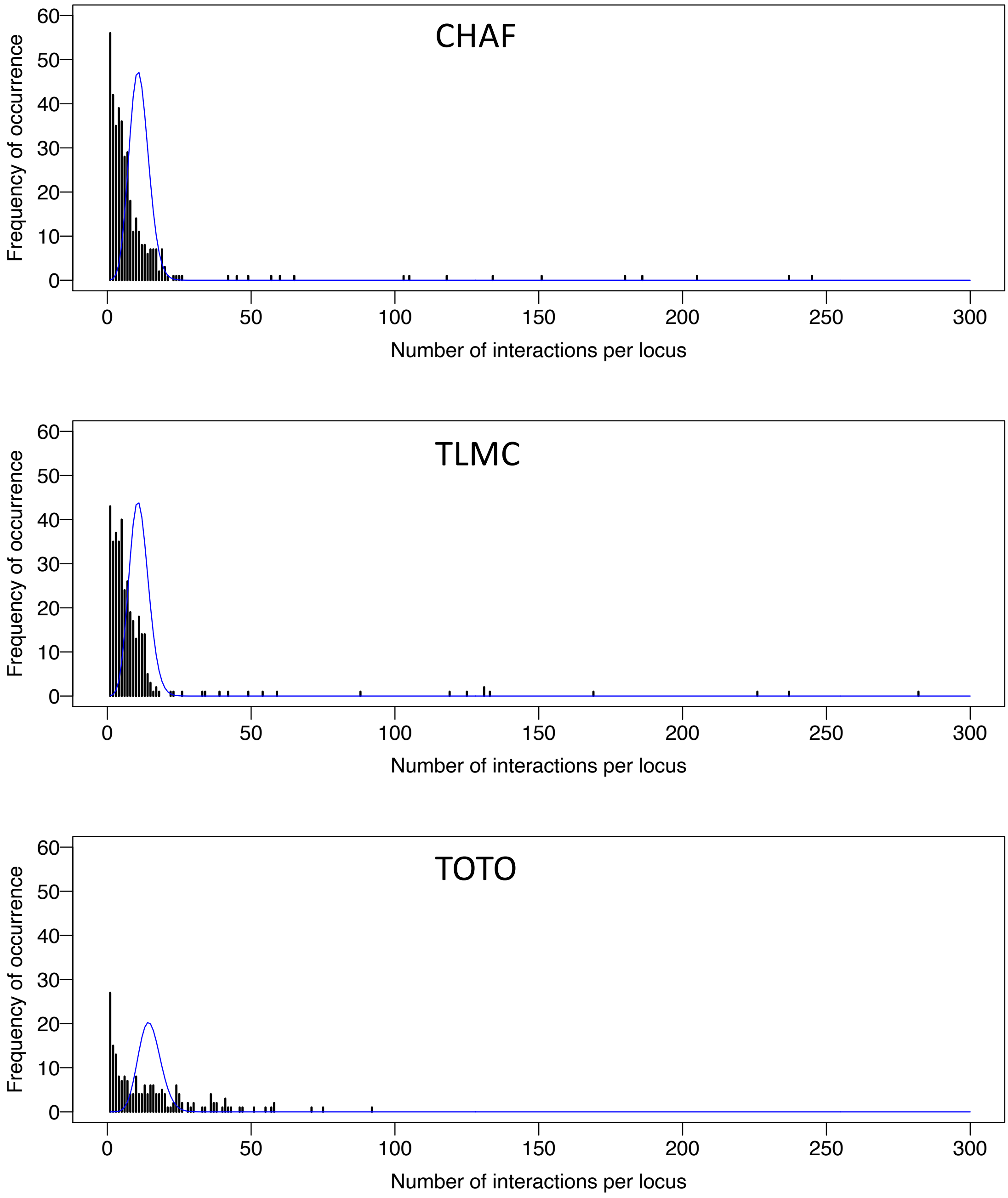


Figure S16. The observed occurrence spectra of detected DMI candidates in the hybrid populations is skewed towards low and high frequencies compared to the expected distribution. The bar plot shows the frequency of occurrence of loci in the putative DMIs for each population. The blue line shows the expected occurrence distribution under a Poisson distribution.

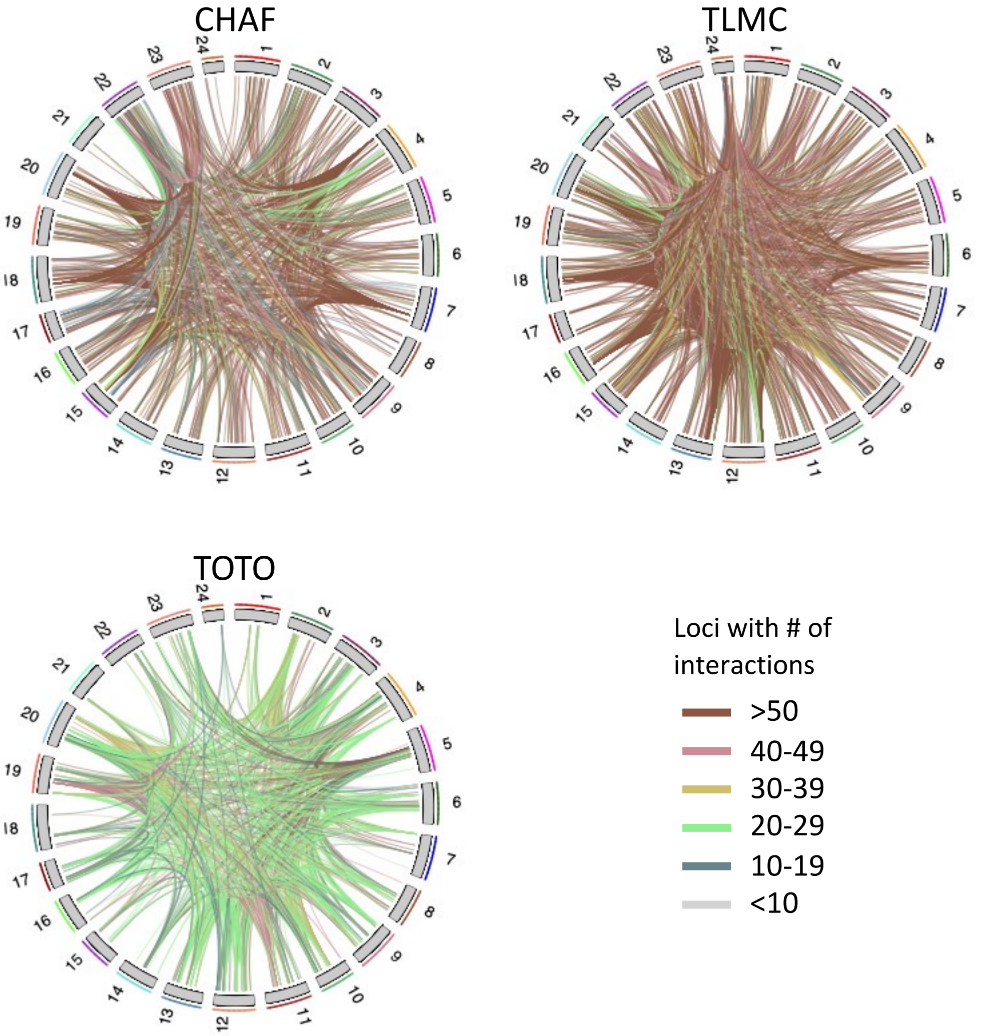


Figure S17. Chromosomal illustration of putative DMIs in three hybrid populations (CHAF, TLMC and TOTO). Some loci have many putative interacting partners. Each line is colored by the number of putative partners of the two involved candidate loci. In TOTO, loci show a smaller number of interactions than in CHAF and TLMC.

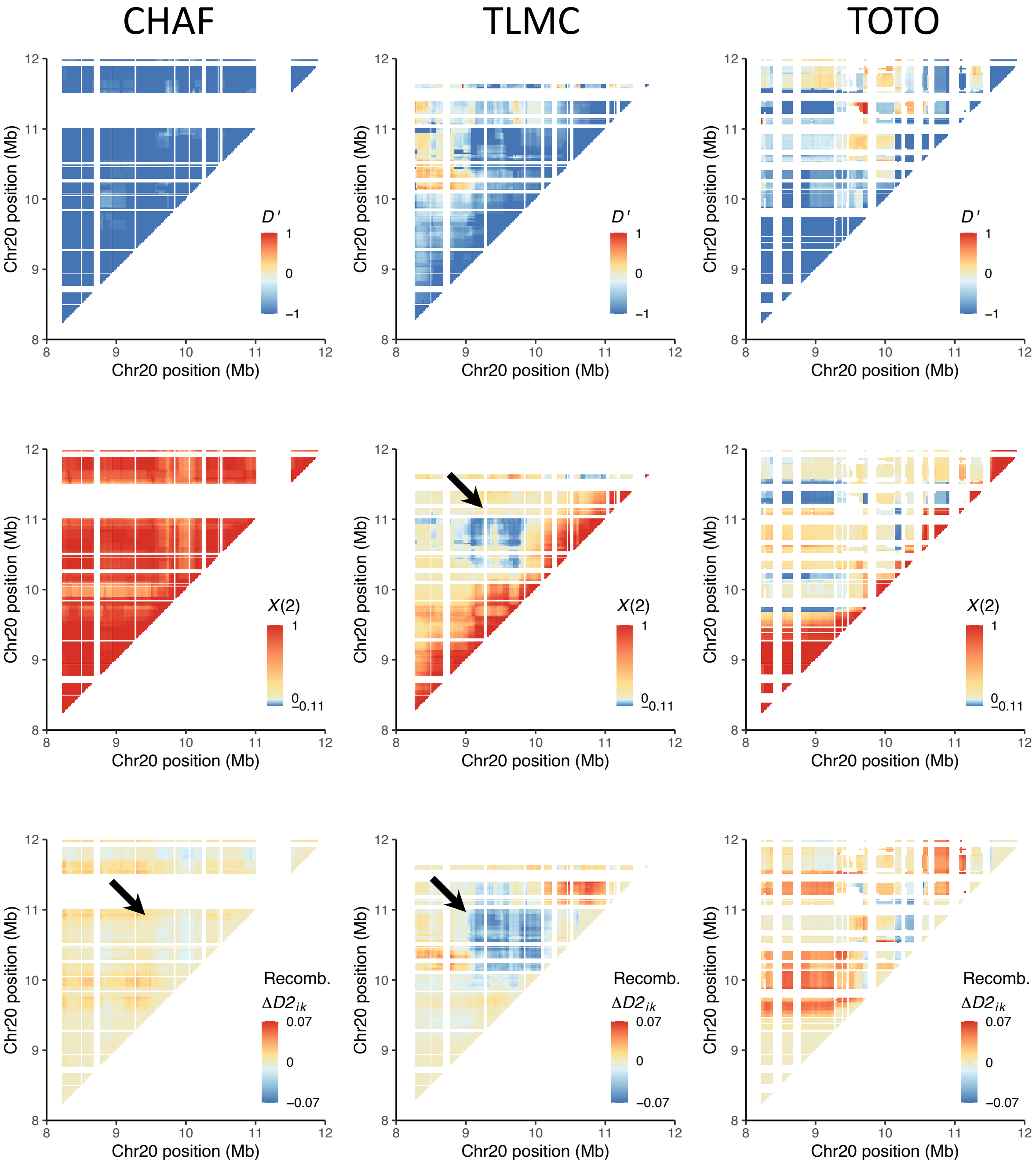


Figure S18. Example of an intrachromosomal DMI candidate in hybrid swordtail fish on chromosome 20. The values of $X(2)$, linkage disequilibrium ($D'$), and recombinant imbalance ($\Delta D2 = D2_{MB} - D2_{BM}$) are presented by color intensity. As highlighted by an arrow, the candidate DMI was detected in TLMC between locus chr20:9Mb (B alleles) and chr20:10Mb (M alleles). In CHAF, the residual parental linkage is still so strong that $X(2)$ >0, but $\Delta D2$ shows a weak recombinant imbalance. In TOTO, a stronger interaction presumably happened between chr20:8Mb and chr20:11Mb such that we see no significant signal between locus chr20:9Mb and chr20:10Mb.

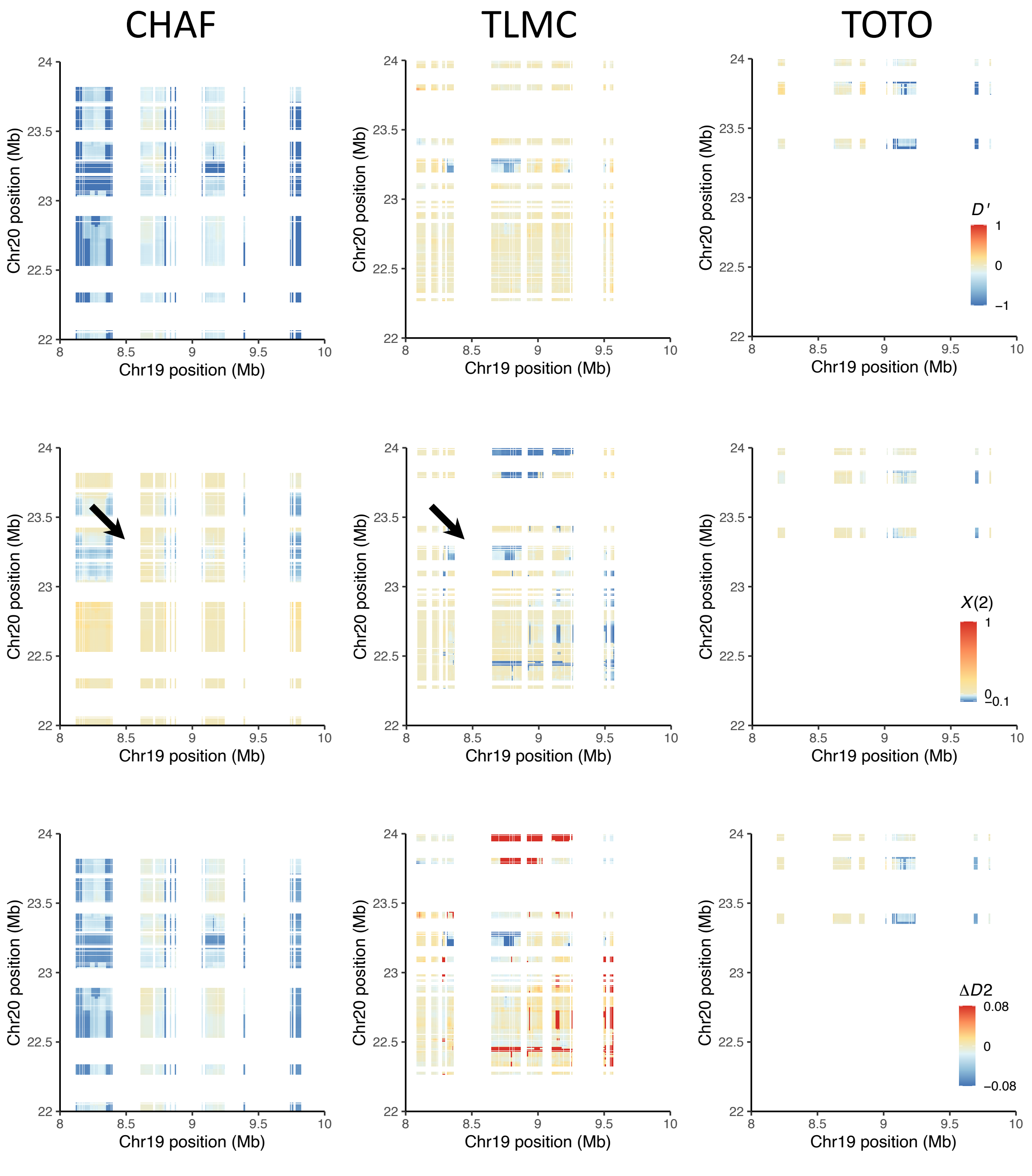


Figure S19. Example of an interchromosomal DMI candidate in hybrid swordtail fish between chromosome 19 and 20. The values of $X(2)$, linkage disequilibrium ($D'$) and recombinant imbalance (Recomb. $\Delta D2 = D2_{MB} - D2_{BM}$) are presented by color intensity. As highlighted by an arrow, the candidate DMI was inferred in TLMC between locus chr19:8-9Mb (B alleles) and chr20:23Mb (M alleles). We suspect that negative $X(2)$ is due to linkage and that the true DMI locus has been fixed in all three populations. Most of the homozygous M alleles at chr20:22-24Mb have been purged in TOTO but not at chr19:8-9Mb. CHAF shows a larger genomic region involved in DMI than TLMC, which might be due to the recombination map at the first several generations after hybridization.

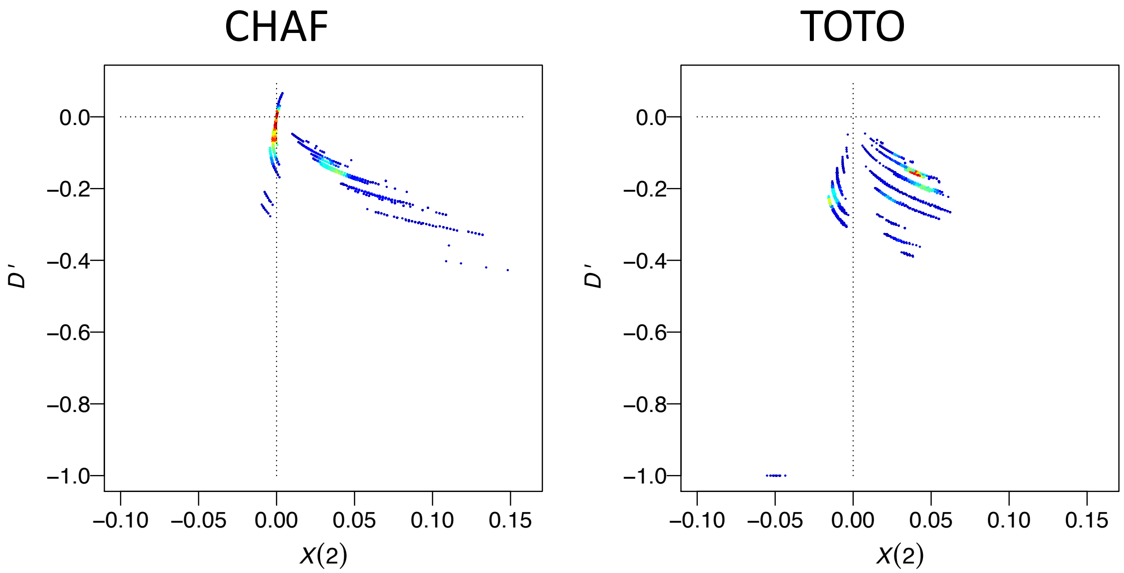


Figure S20. Scatterplot of $X(2)$ and $D'$ for markers occurring at *Xmrk* and its interacting locus *cdkn2a/b*. Markers were extracted from the focal genes and their flanking regions (*cdkn2a/b at* chr5:15.7Mb-16Mb*, Xmrk* at chr21:16.5Mb-17.5Mb). No pairs of markers in TLMC passed the filtering criteria. A large proportion of pairs are $D' < 0$ & $X(2) < 0$ in both CHAF and TOTO between *Xmrk* and *cdkn2a/b.* Negative $X(2)$ values are larger in TOTO (age, ~35 generations) than in CHAF (age, ~25 generations). In TOTO, $D' = -1$ & $X(2) < 0$ indicates the absence of genotypes that are linked to DMIs, possibly due to sampling at low frequencies or completed purging of incompatible alleles. Colouring indicates point density.

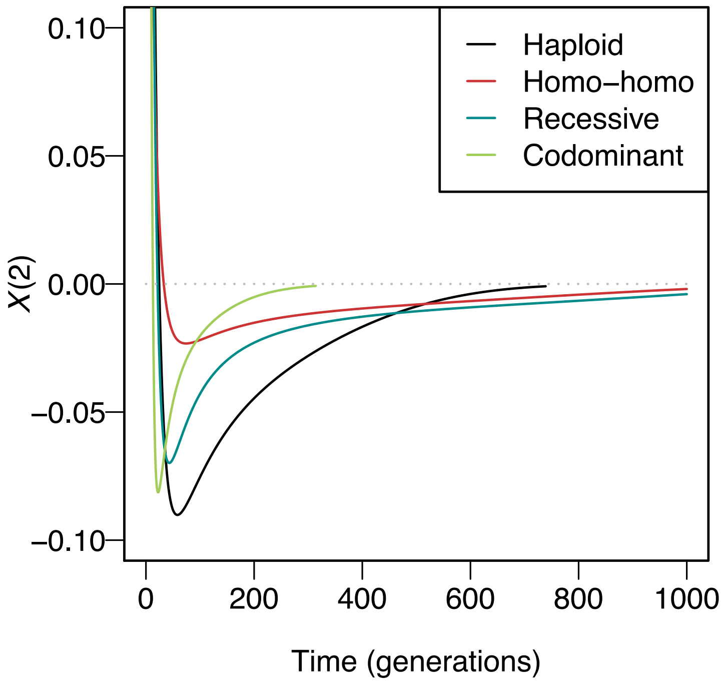


Figure S21. Dominance effects on $X(2)$ dynamics of DMIs. The four DMI scenarios are a DMI in the haploid model, homozygous-homozygous DMI (Homo-homo), homozygous-heterozygous DMI (recessive), and heterozygous-heterozygous DMI (codominant). All trajectories were generated with the same parameters. Incompatible alleles $A$ and $B$ are under weak direct selection ($\alpha = 0.001$ for $A$ and $\beta = 0.002$ for $B$). The strength of the epistatic interaction between alleles $A$ and $B$, $\gamma$, is -0.5. The recombination probability $c$ is 0.1. The admixture proportion is 0.5.

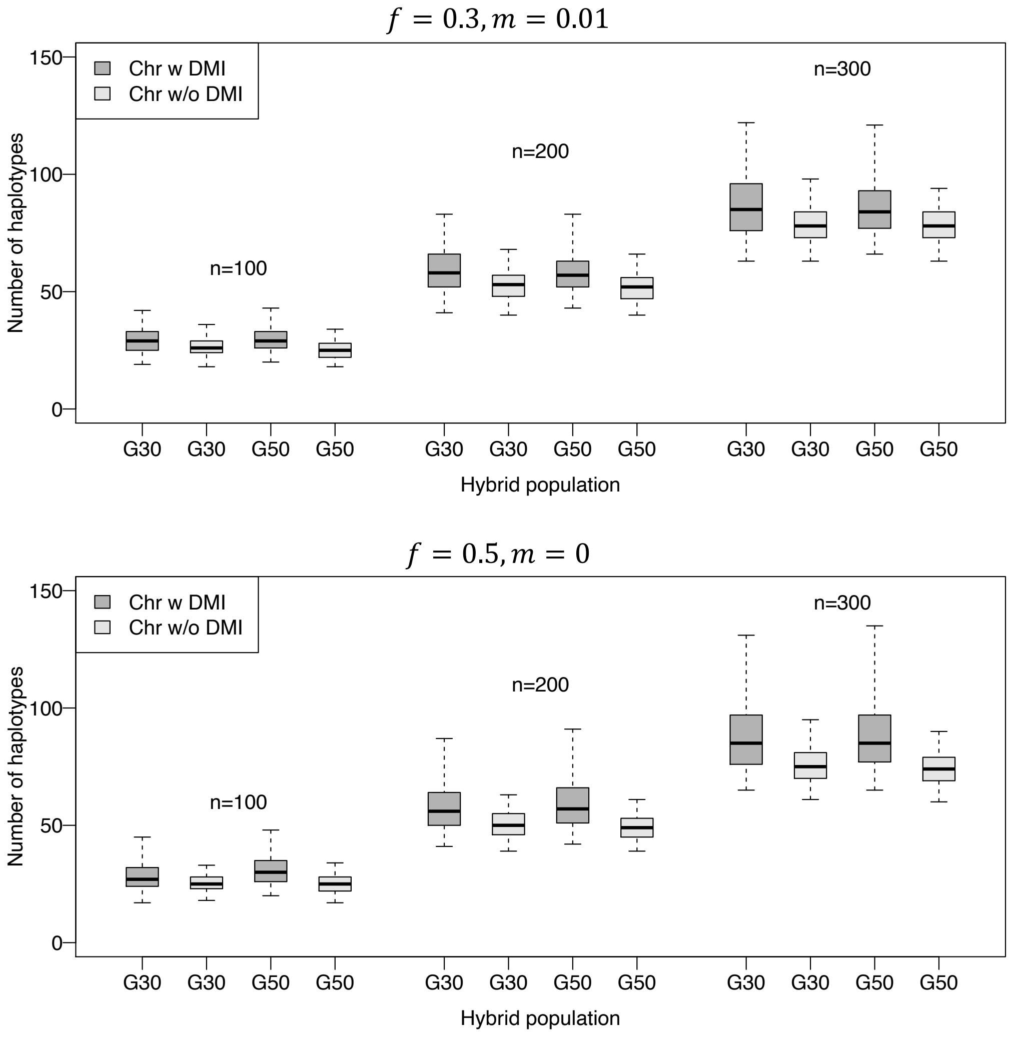


Figure S22. The number of two-locus homozygous-homozygous haplotypes for different sampling sizes ($n = 100, 200, 300$), obtained from a set of 100 simulations for each parameter combination using SLiM. Simulations were performed assuming a diploid Wright-Fisher population with 5000 individuals. In each simulation, two DMIs were randomly distributed on four chromosomes (Chr w DMI). Two additional neutral chromosomes (Chr w/o DMI) were also simulated in the same genome. The number of two-locus homozygous-homozygous haplotypes for each pair loci were counted at generation 30 and 50 (G30 & G50). Neutral chromosomes contain fewer two-locus homozygous-homozygous haplotypes on average than chromosomes with DMIs. Boxes represent the interquartile range, and whiskers extend to 95% confidence interval.
